# A scalp geometry based parameter-space for optimization and implementation of conventional TMS coil placement

**DOI:** 10.1101/2022.01.22.477370

**Authors:** Yihan Jiang, Boqi Du, Yuanyuan Chen, Lijiang Wei, ZhengCao Cao, Zong Zhang, Cong Xie, Quanqun Li, Zhongxuan Cai, Zheng Li, Chaozhe Zhu

## Abstract

Transcranial magnetic stimulation efficacy is largely dependent upon coil position and orientation. A good method for describing coil placement is required for both computational optimization (planning) and actual placement (implementation). In coordinate dependent parameter-spaces (CDPs), three-dimensional coordinates are used to represent coil position and three orthogonal unit vectors are used to represent coil orientation. A CDP can precisely describe arbitrary coil placement; therefore it offers great advantage in computational optimization which checks through all possible placements. However, a neuronavigation system is usually required to accurately implement the optimized CDP parameters on a participant’s head. Routine clinical practice, on the other hand, often uses the International 10-20 system to describe coil placement. Although the 10-20 system can only perform modeling and placement at limited scalp landmarks, it allows the synthesis of different individuals’ targeting effects to find group-optimal parameters; it also allows manual placement, which is important for commonly-seen use cases without individual MRI scans and navigation devices. This study proposes a new scalp geometry based parameter-space (SGP), integrating the advantages of CDP and 10-20 methods. Our SGP 1) can quantitatively specify all possible conventional coil positions and orientations on an individual’s scalp, which is important for electrical modeling and optimization, 2) maintains inter-individual correspondence, which is important for synthesizing TMS effects from different individuals and studies. 3) enables fast and simple manual implementation. Demonstration experiments were conducted to illustrate the application of an SGP-based framework for both individual and group-based optimization. A measurement experiment was performed to evaluate speed, precision and reliability of SGP-based manual implementation; results show it surpasses previous manual placement methods.

## 1. Introduction

Transcranial magnetic stimulation (TMS), one of the most important in vivo neuromodulation techniques in both basic and clinical neuroscience (Lefaucheur et al., 2020), can cause instant or long-term changes in cognition and behavior by inducing electric currents in the brain via rapidly changing magnetic fields generated by a stimulating coil placed on the scalp (AT et al., 1985). The importance of both coil location and orientation to TMS efficacy have been well demonstrated. For example, it has been reported that TMS antidepressant efficacy is better in lateral and anterior prefrontal coil locations, where underlying cortical regions are more anti-correlated to the subgenual cingulate (Fox et al., 2012; Herbsman et al., 2009). The amplitude of TMS motor-evoked potential (MEP) can vary markedly due to small coil orientation changes of just 10° to 15° (Tarapore et al., 2013). Such coil position and orientation dependent effects have also been investigated in computational modeling studies, which have found that coil position and orientation have complex interaction effects with brain structure, leading to different strengths of induced electric fields. Therefore, coil position and orientation on the scalp should be optimized to ensure efficient stimulation of the target region (Gomez et al., 2021).

Coil placement optimization usually consists of two stages: planning and implementation. In the planning stage, the electric fields induced by all possible TMS coil placements are simulated based on an individual’s brain structure. This process finds the optimal placement parameters, those that maximize the chosen effect index, for example, the E-field strength at the cortical region of interest (ROI) (Balderston et al., 2020). An electric-field-modeling-based optimization framework for coil placement has been proposed and validated with physiological observations (Opitz et al., 2013; Weise et al., 2020). A fast computational auxiliary dipole method as well as deep-learning-based electric field modeling have further enabled exhaustive search of parameters (Gomez et al., 2021). However, individualized optimization is not a common in routine TMS treatment, due to the need for MRI scans and to the time, computing power and technical proficiency with a complex image analysis pipeline necessary. Therefore, synthesizing individual optimization results into a group-based coil placement atlas would be an important advance, one which can support the coil placement decision process during typical interventions, especially large-throughput clinical practice (Gomez-Tames et al., 2018a, 2020). In the implementation stage, the coil should be placed on the individual’s head according to the optimized parameters. The most precise method is using a stereotaxic neuronavigation system to guide placement. However, such an expensive device is unaffordable in many clinical settings, and placements based on manual measurement have become an important alternative (Beam et al., 2009; Vaghefi et al., 2015).

Appropriately describing the position and orientation of a coil placement is the common basis of both simulation and optimization *in silico* and actual coil placement *in vivo*. Currently, there are two main types of coil placement descriptions. One type is based on the International 10-20 EEG reference system and its derivatives (10-10 and 10-5 systems), while the other type is based on three-dimensional coordinate systems. The International 10-20 system (JASPER & H., 1958) is a proportional scalp landmark system, consisting of 25 landmarks, 4 initial reference points and 21 evenly distributed scalp points iteratively defined by relative proportional distances between prior landmarks, thus taking inter-individual variations in head size into account. The 10-20 system was the first coil location description method and has been widely used, such as in the well-known F3 treatment for depression. When using this method, the coil is usually placed tangent to the scalp, with coil center contacting the scalp (called conventional placement) to ensure minimal energy attenuation and easy operation. The coil position (i.e., coil center location) is described by one 10-20 point and the coil orientation (i.e., coil handle direction) can be described using an additional 10-20 point to specify the direction the handle points (Saturnino et al., 2019), or more commonly, using the degree between the handle and the mid-sagittal plane (Lefaucheur et al., 2020). The 10-20-system-based description intrinsically provides inter-individual consistency, allowing optimization results from different individuals to be directly combined to form a group-based optimization atlas (Gomez-Tames et al., 2020). However, the 10-20 system includes only 25 scalp positions, so the describable placement space is highly limited. The 10-10 and 10-5 systems provide higher spatial resolution, but they are still enumeration methods, rather than continuous coordinate systems, and therefore unable to describe all possible placements. This may mean that the optimal coil placement is missed. When considering the measurement of points on the scalp, each 10-20 system point can theoretically be measured manually, but the identification of 10-20 landmarks in later steps depends on the positions of 10-20 landmarks determined in prior steps (JASPER & H., 1958; Milnik, 2009), making this procedure time-consuming (16 min reported by (Xiao et al., 2017)) and error prone. The Beam F3 method (Beam et al., 2009) and a semi-automatic 10-20 navigation system (Xiao et al., 2017) have been proposed to improve the speed and reliability of 10-20 measurements.

Coordinate dependent parameter-spaces (CDPs) are more powerful descriptive methods for coil placement. CDPs use three-dimensional coordinates and three orthogonal unit vectors to represent position and attitude of the coil (Saturnino et al., 2019). Unlike the 10-20 system, a CDP is a continuous space that can quantitatively describe arbitrary coil placement, which is important for global optimization. One problem is that this space includes regions inside the person’s head (i.e., the coil intersects with the head), which are physically inaccessible, as well as large regions far away from the head, where coils are unable to generate efficient stimulation. Therefore, most optimization studies have used conventional placement spaces as search spaces, to reduce the size of the search space and improve the efficiency of optimization results (Balderston et al., 2020; Gomez et al., 2021; Gomez-Tames et al., 2018b). Unlike the 10-20 system, optimal CDP parameters of different individuals are difficult to synthesize. Individually optimal CDP parameters come from different 3D coordinate systems based on different MRI spaces, with different head size and shape. Therefore, how to summarize CDP optimization results from different individuals is a complex problem. In addition, the optimal CDP placement parameters (coil position coordinates and orientation vectors) cannot be directly manually measured. They instead require interactive navigation based on neuronavigation systems for implementation. This is also a reason why CDP is not used more commonly.

Taken together, 10-20-based methods and CDP-based methods each have their own advantages in individualized optimization, group synthesis and manual placement, but also drawbacks. The present study aims to combine the advantages together: we propose a scalp geometry based parameter-space (SGP) that can 1) quantitatively specify all possible positions and orientations in conventional placement; 2) maintain inter-individual comparability; 3) be implemented *in vivo* in a fast and simple way, with only tape measure and pen. In the present study, we first provide the SGP definition, then we present a general framework for optimization and implementation of SGP-based coil placement, including both individualized and non-individualized optimization. We provide a demonstration experiment to give a detailed and intuitive presentation of how to apply SGP-based systems during individualized and group-based optimization of TMS coil placement. For the implementation stage, we specifically propose a manual placement protocol that can easily be implemented for arbitrary positions and orientations described by SGPs. We perform two preliminary experiments to evaluate the speed, precision and reliability of manual placement.

## 2. Methods

### 2.1 SGP space (*s, φ*) for conventional TMS coil placement

Like most individual coil placement optimization studies (Balderston et al., 2020; Gomez et al., 2021; Gomez-Tames et al., 2018b), we also perform optimization in the conventional placement space, defining the coil position as the location of the contact point between the coil center and the person’s head and defining the coil orientation as the angle of the handle (in the plane tangent to the head surface) versus the mid-sagittal plane.

We utilize our previously proposed Continuous Proportional Coordinate (CPC) system to define the position parameters of an arbitrary scalp point *s* (P_NZ_, P_AL_) (Fig. 1A). Like the 10-20 system, CPC parameters are calculated from head reference points: nasion (NZ), inion (IZ) and the left/right preauricular points (AL/AR). Given these reference points, the CZ point, also needed for the procedure, can be identified as follows. First, find the midpoint of an arbitrary NZ-IZ curve, Mid_NZIZ_. Next, find the midpoint (called Mid_ALAR_) of the curve going through AL, Mid_NZIZ_ and AR. Next, find the midpoint of the curve going through NZ, Mid_ALAR_ and IZ, giving a new Mid_NZIZ_. Repeat this process until the iteratively found Mid_NZIZ_ meets the iteratively found Mid_ALAR_: this intersection midpoint is CZ.

**Figure 1.**
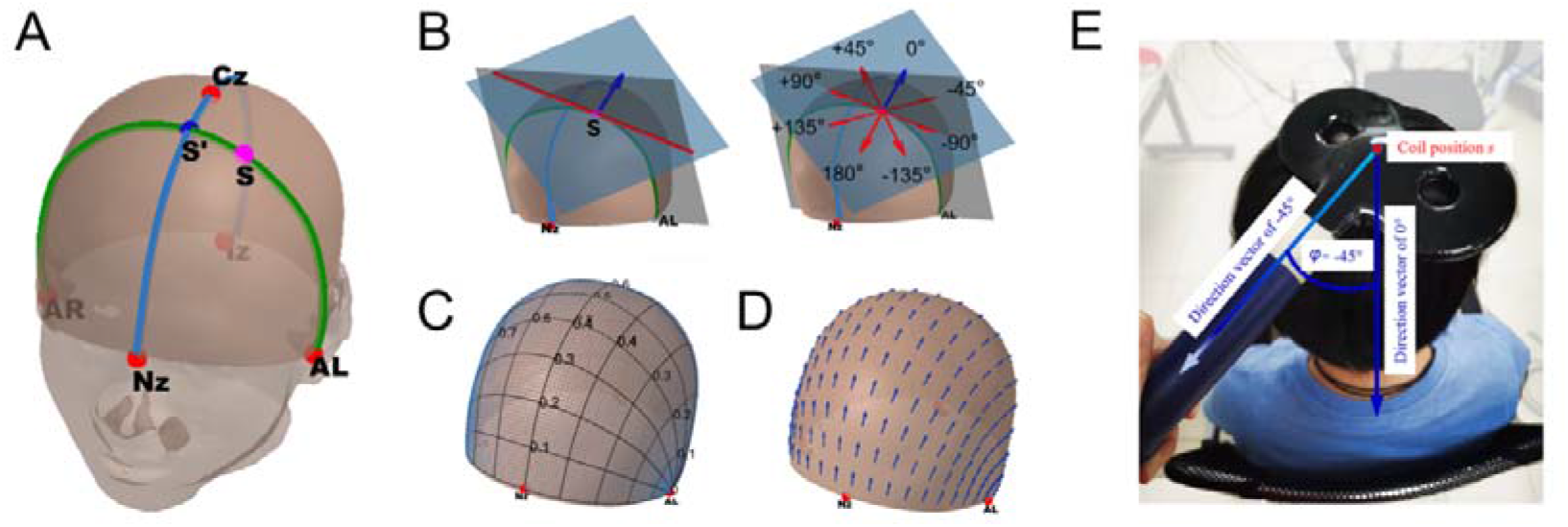
(A) Definition of coil an example position *s* (magenta dot, P_NZ_ = 0.4, P_AL_ = 0.4) involves proportions of lengths along two curves anchored at cranial landmarks. (B) Definition of zero-degree direction (blue vector) and example direction vectors of different orientation *φ* (red vectors) at coil position *s*. (C) A discrete version of SGP obtained by evenly dividing each dimension into 10 units. (D) Zero-degree direction vectors at different coil positions. (E) An example showing placement of a coil at position *s* with orientation *φ*= -45°.

Then, the coordinates of any point *s* can be determined by proportional distances on two geodesic curves. One is a reference curve that is independent of *s*, defined as the intersection of the scalp surface and the plane through NZ, CZ and IZ (i.e., mid-sagittal plane; blue curve in Fig. 1A). The length of the reference curve is denoted as L_ref_. The other curve, the *active* curve, depends on *s* and is defined as the intersection of the scalp surface and the plane through AL, AR and *s* (green curve in Fig. 1A). The length of the active curve is denoted as L_active_. The reference curve and active curve intersect at *s*’. The first position parameter P_NZ_ is the proportion between curve length from NZ to *s*’ and the full reference curve length, i.e., P_NZ_ = L_NZ-s’_/L_ref._ The second position parameter P_AL_ is calculated as the proportion between curve length from AL to *s* and the full active curve length, i.e., P_AL_ = L_AL-s_/L_active_. In this way, any scalp point *s* has a pair of corresponding position parameters (Fig. 1B).

After defining coil location *s*, the orientation can be defined in the tangent plane of point *s* (Fig. 1B). We first specify a zero-degree direction. The gray plane (Fig. 1B) is determined by AL, AR and *s*, while the blue plane is the tangent plane at point *s*. These two planes intersect at the red line. The zero-degree direction is defined by a unit vector in the tangent plane that is perpendicular to the red line, starting at *s* and pointing backwards (blue vector). Any orientation can be obtained by rotation of the zero-degree direction vector (positive anticlockwise, negative clockwise) in the tangent plane. Taking the figure-8 coil as an example, the SGP parameters of a coil placement is shown in Fig. 1E.

### 2.2 SGP based optimization and implementation of coil placement

We first show the pipeline when conditions for individual optimization are available (individual MRI scan provided, computing power and time are sufficient). At the planning stage, the individual’s T1 image is segmented into different issues to generate the individual’s head mesh and scalp mesh. Using the individual’s scalp mesh, search space range and search interval as input, a discrete SGP search space {(*s*_k_, *φ*_q_), k=1…, K; q=1…, Q} can be constructed. By changing total position number K and total orientation number Q, the search space can be defined with arbitrary density.

**Figure 2.**
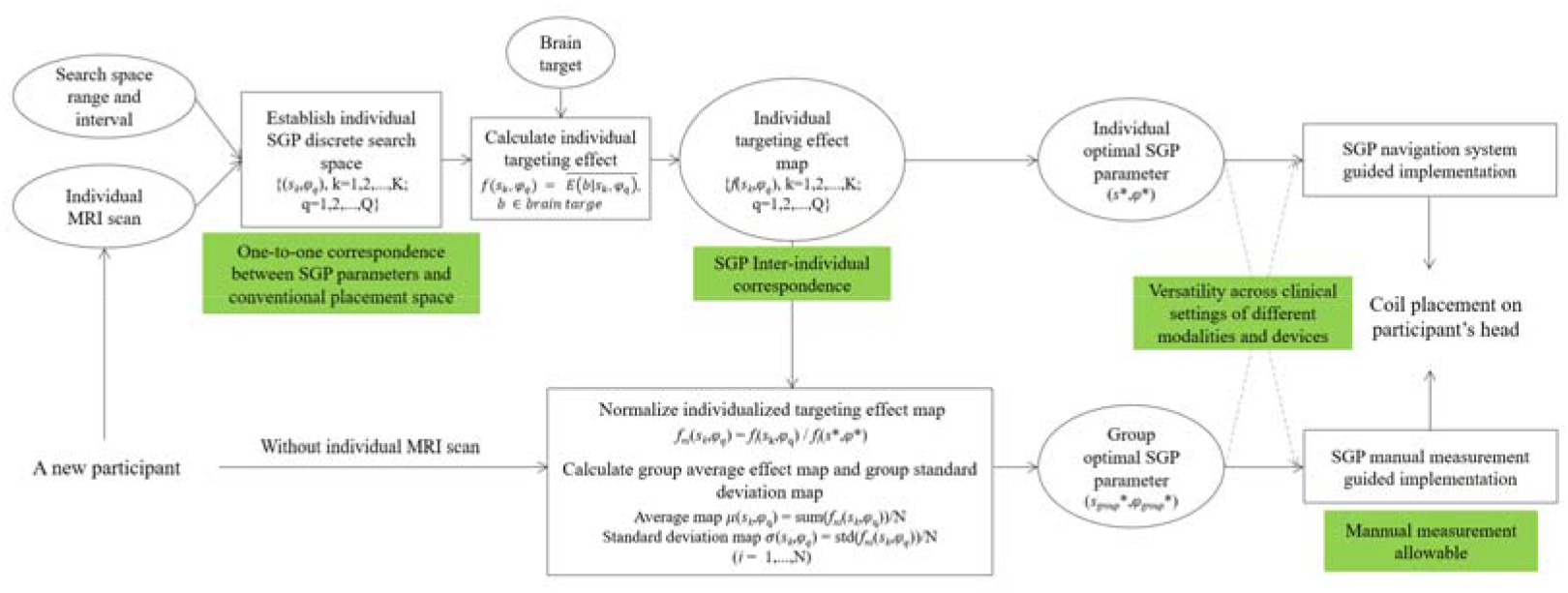
Framework for SGP-based optimization and implementation of coil placement. Individualized optimization pipeline is shown at the top, and group-based optimization pipeline without individual MRI data is shown at the bottom. In the last step, solid lines point to the typical implementation, while dotted lines point to alternatives. The main properties of SGP that enable the process are emphasized in green boxes.

We can then simulate the electric field distribution E(*s_k_, φ_q_*) for each placement in the search space based on the head model mesh constructed from the individual’s data. Additionally, stimulation intensity, coil configuration and tissue electrical conductivities can be added into this modeling process, or default parameters (provided in software like SimNIBS (SimNIBS Developers, 2020)) can be used. Next, given a brain target (can be defined using anatomical, functional or connectivity properties) and an optimization strategy (here e.g., maximize the electric field strength inside the target), we can calculate the targeting effect *f*(*s_k_,φ_q_*) as the average electric field strength inside the target for each placement (*s_k_,φ_q_*) and find the optimal parameter (*s_*_,φ_*_*) for this individual:

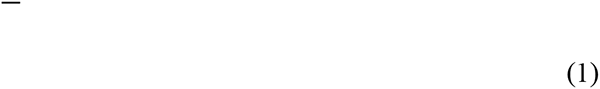

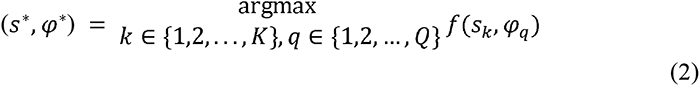

At the implementation stage, the coil placement can be implemented on the person’s head very precisely with the guidance of an SGP-based navigation system (in development). Alternatively, placement can be indirectly implemented by transforming the SGP parameters to three-dimensional CDP parameters (when the individual’s T1 image is available) and using a CDP neuronavigation system (like BrainSight) to guide placement. Another alternative is to implement placement via manual measurement.

In circumstances not permitting individualized optimization, one possible solution is to summarize the normalized targeting effect map *f_ni_* (*s_k_,φ_q_*) from different individuals *i* ( *i* =1,2,…,N) in a database into a group average targeting effect map to guide placement. SGP parameters intrinsically correspond across individuals, therefore, if the search space is the same for different individuals, the results can be directly synthesized.

First, for each individual *i*, the individual’s targeting effect at (*s_k_,φ_q_*) are normalized by dividing by the maximum targeting effect.

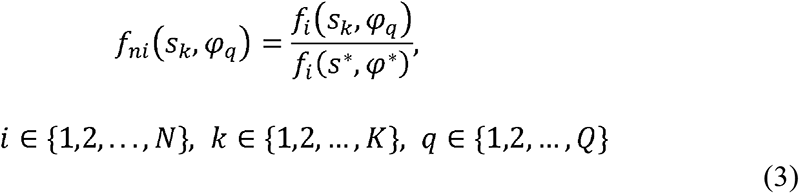

Then, the normalized individual targeting effects of many individuals can be synthesized into an average targeting effect map.

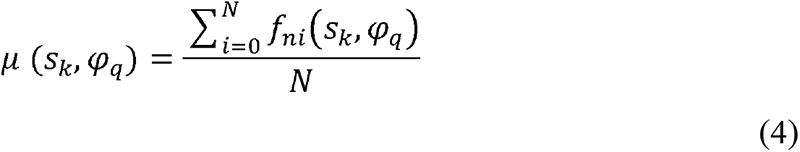

The standard deviation of the targeting effect map across different individuals can also be calculated.

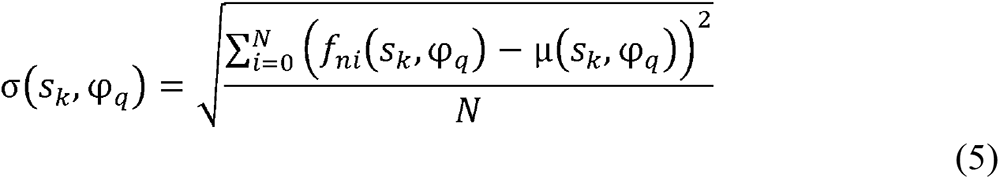

If there exists a placement parameter setting with relatively large average targeting effect *μ* and relatively small standard deviation *σ*, then it can be chosen as a group-appropriate placement parameter (*s_group_^*^, s_group_*φ*^*^*) . The group-appropriate placement can be implemented through an SGP-based navigation system or through SGP-based manual measurement.

### 2.3 SGP Manual measurement

Tools needed in the manual measurement process include two tape measures, a marker, a protractor (or visual estimation) and a calculator (or mental calculation).

For any given position *s* (P_NZ_,P_AL_) and orientation *φ*, the measurement steps are as follows:

1. Find 4 reference points, the Nasion, Inion and left and right preauricular points on the head (Fig. 3[1]). Time: about 30 seconds.
2. Find point the CZ point and the reference curve. Time: about 3 minutes. First, find the midpoint of an arbitrary NZ-IZ curve, Mid_NZIZ_. Next, find the midpoint of the curve connecting AL, Mid_NZIZ_ and AR, Mid_ALAR_. Next, find the midpoint of the curve connecting NZ, Mid_ALAR_ and IZ and set it as a new Mid_NZIZ_. Iterate the above two steps until Mid_NZIZ_ meets Mid_ALAR._ This common midpoint is CZ. The curve connecting NZ-CZ-IZ is the reference curve (Fig. 3[2]).
3. Find point *s*’, the active curve and point *s*. Time: about 1.3 minutes First, measure the length of reference curve L_ref_, calculate the length value P_NZ_×L_ref_. Then, mark point *s*’ as the point along the reference curve that is P_NZ_×L_ref_ distance from NZ. The curve connecting AL, *s*’ and AR is the active curve. Next, measure the length of active curve L_active_ and calculate the length value P_AL_×L_active_. Then, mark point *s* as the point along the active curve that is P_AL_×L_active_ distance from AL (Fig. 3[3]).
4. Find the -90° direction and arbitrary orientation *φ*. Time: about 20 seconds. The direction from point *s* to AL along the active curve is the -90° direction. Using a protractor (or visual estimation), find orientation *φ* based on the -90° direction ([Fig. 3[4]).
5. Use a marker or an eyebrow pencil to mark point s and orientation *φ* on the cap (Fig. 3[5]). Time: about 40 seconds.

**Figure 3.**
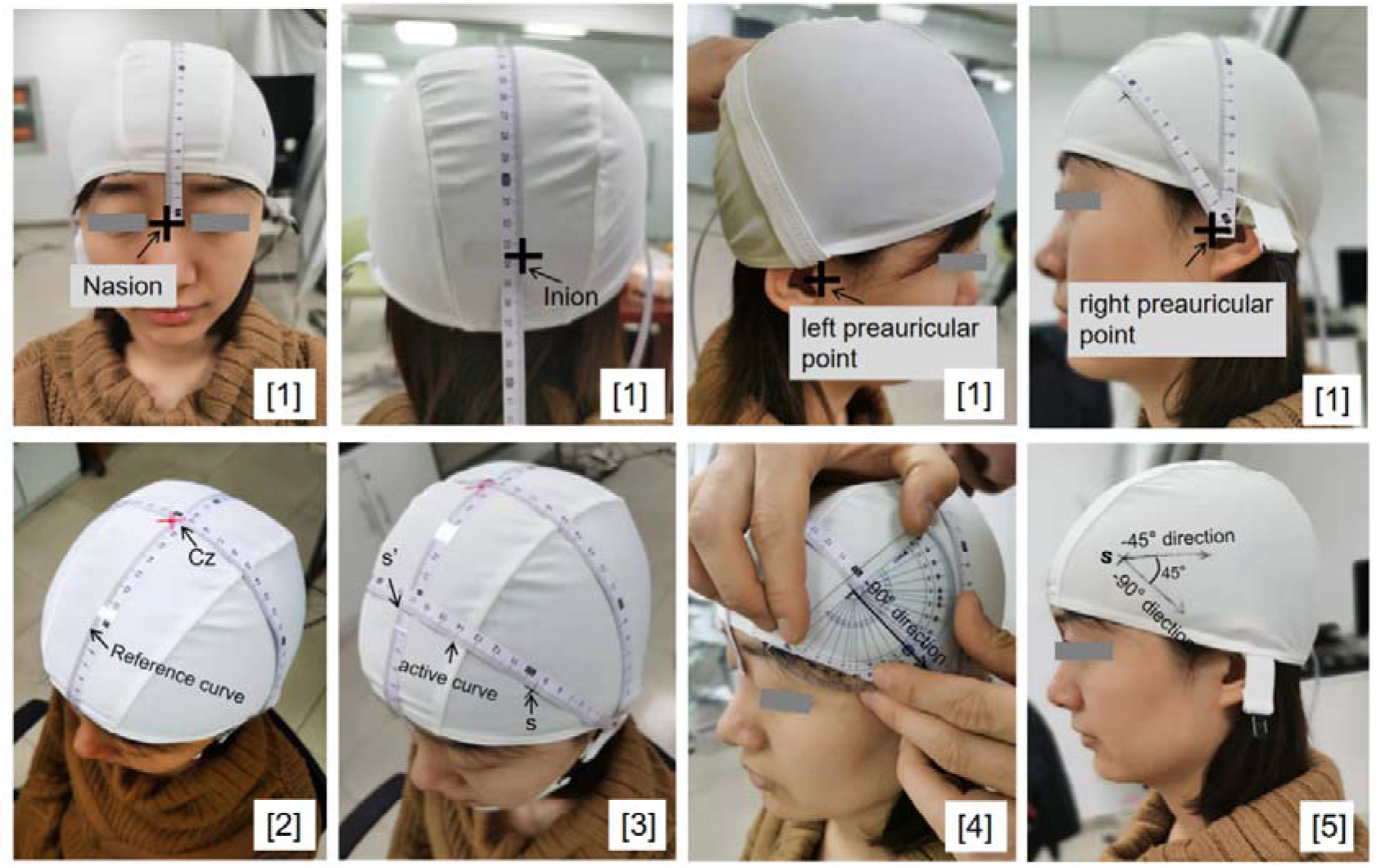
Illustration of manual measurement of SGP parameters. (1) Mark 4 reference points: Nasion, Inion and left and right preauricular. (2) Find CZ and the reference curve. (3) Find point s’, the active curve and point s. (4) Find the -90° direction and orientation *φ*. (5) Mark point s and orientation *φ* on the cap.

### 2.4 Manual measurement experiment: evaluating speed, precision and reliability

Five healthy adults (mean age = 23.7, 3 females, 2 male) were enrolled in a preliminary study to investigate the speed, precision and reliability of manual position measurement. Nine typical positions (P_NZ_, P_AL_: (0.27,0.26), (0.53,0.25), (0.79,0.24), (0.22,0.47), (0.52,0.50), (0.78,0.53), (0.24,0.72), (0.50,0.77), (0.75,0.75)) uniformly distributed on the scalp surface were set as targets. For each participant, nine positions were measured by three trained TMS technicians at two time points over the course of one month, with at least 24 hours separating measurement sessions (n = 270 total measurements, 30 at each scalp target). The first step of position measurement, finding the CZ and reference curve, is the same for different targets and therefore that process was performed only once at the start and shared for other points.

We used a commercial 3D digitizer (Fastrak™, Polhemus) to perform a dense sampling of each participant’s head from which the head point cloud was reconstructed. The position of the point identified by manual measurement was recorded by the 3D digitizer. The position of the target point was calculated on the reconstructed head point cloud (acting as ground truth). Distance between the identified point and ground truth target point was calculated as the measurement error (quantifying precision). We separately evaluated inter-technician reliability relative to a group-averaged target coordinate and intra-technician reliability relative to a technician-specific average (following the reliability definition in (Trapp et al., 2020)).

Furthermore, four healthy adults (mean age = 25.3, 4 females, 1 male) were enrolled in a preliminary study to investigate the precision of the manual orientation measurement. For each participant, four orientations (0°, 45°, 90°, 135°) were measured at three scalp positions ((0.27,0.26), (0.52,0.5), (0.75,0.75)) by three trained TMS technicians (n=144 total measurements), following the manual measurement protocol. Additionally, technicians also measured 45° via visual observation, to evaluate the precision of this common clinical practice. The direction vector of the manually identified orientation was recorded by the 3D digitizer; the direction vector of the target orientation was calculated from the participant’s head point cloud generated by dense sampling on the head (acting as ground truth). The angle between the identified orientation and the ground truth target orientation was calculated as the measurement error (quantifying precision).

### 2.5 Demonstration experiment for individualized and group-based optimization

A demonstration experiment using real data was conducted to illustrate how the SGP-based optimization protocol works. We show the utility of this protocol in generating “coil placement targeting effect atlases” that relate coil placement to individualized or group-based targeting effects. These atlases can be used to guide researchers and clinicians in deciding optimal coil placement for a specific brain target. The brain target in this demonstration is the motor cortex.

#### Input MRI Data Acquisition

Each individual’s high-resolution T1-weighted structural image was acquired on a Siemens Trio 3T MRI Scanner (TR/TE, 2530/3.5ms; flip angle, 9°; field of view, 176 mm × 256 mm; slices, 256; thickness, 1.0 mm; voxel size, 1 mm × 1 mm × 1 mm). The individual’s BOLD functional images were acquired on a Siemens Trio 3T MRI Scanner (TR/TE, 2000/28ms; flip angle, 90°; field of view, 102 mm × 102 mm; slices, 32; thickness, 2.0 mm; voxel size, 2 mm × 2 mm × 2 mm; volumes, 156) during a finger tapping task, including seven rest blocks of 24 s with fixation point and six task blocks of 24s with right-hand tapping by the first dorsal interosseous muscle. Four dummy scans were completed at the beginning of each run to allow for stabilization of the MR signal. To achieve a higher resolution, BOLD images were focused to cover the cortical motor areas, so an additional whole EPI volume was acquired for co-registration (TR/TE, 6000/28ms; flip angle, 90°; field of view, 102 mm × 102 mm; 96 slices; 2 mm isotropic resolution).

#### Experimental procedure

1. Based on the SPM12 (https://www.fil.ion.ucl.ac.uk/spm/software/spm12/, Ashburner and Friston, 2005) segmentation routine, the individual’s T1 image was segmented into six tissue images (gray matter, white matter, cerebrospinal fluid (CSF), bone, soft tissue, and air/ background). Subsequently, the scalp mesh was extracted from a smoothed and binarized head image (gray matter + white matter + CSF + bone + soft tissue) (Xiao et al., 2018) and the head model mesh was created with SimNIBS’s default pipeline “headreco”(Nielsen et al., 2018). <u>Time: about 2 hours.</u>
2. We identified four cranial landmarks NZ, AL, AR and IZ in the individual T1 image using MRIcron software and located the CZ point using Jurcak’s iterative algorithm (Tsuzuki et al., 2007). Based on these reference points, a discrete SGP space was established by uniformly dividing the geodesic curves (including active curve and reference curve) into 100 portions (spatial resolution = 1%, i.e., ΔP_NZ_ = 1% reference curve, ΔP_AL_ = 1% active curve) and the angle into 8 intervals (Δ*φ*= 45°). The search space was sampled from this discrete SGP space. The search had 31×31 grid positions (P_NZ*k*_ = 0.3: 0.01: 0.6, P_AL*k*_ = 0.2:0.01:0.5) centered around the scalp point (P_NZ_ = 0.5, P_AL_ = 0.35), which is supposed to be the best target for the motor system according to a meta-analysis study (Jiang et al., 2020)). The inter-point distance was 1% of the reference or active curve length. Four coil orientations were searched per position: from 0° to 135° at 45° intervals. 961 coil positions and 4 orientations combined into 3844 coil placement configurations {(*s_k_,φ_q_*) k=1,2,…,961 q=1,2,…,4}, as shown in Fig. 4.
3. For each coil placement configuration, a three-dimensional scalp coordinate corresponding to the coil position *s_k_* and three three-dimensional vectors X, Y, Z corresponding to the coil orientation *φ_k_* were obtained on the scalp mesh using custom Python code. Specifically, the X vector of SimNIBS corresponded to the coil handle direction vector in our definition, the Y vector corresponded to the normal vector to the tangent plane, and the Z vector was the cross product of X and Y. Given the converted three-dimensional coil placement parameters and the head mesh model as input, we ran simulation in SimNIBS to calculate the induced E-field distribution E(b|s*_k_,φ_q_*). The electrical conductivities (S/m) were set as the defaults, white matter: 0.126, gray matter: 0.275, CSF: 1.654, bone: 0.01, scalp: 0.465, eyes: 0.5, silicone rubber: 29.4 and saline: 1.0. After the electric-field modeling, the targeting effect *f*(*s*_*k*_,*φ*_*q*_) was calculated for each (*s*_*k*_,*φ*_*q*_). Following the protocol proposed by Balderston (2020), we defined *f*(*s*_*k*_,*φ*_*q*_) as the electric field strength inside an individualized region of interest (ROI). The individualized motor ROI was set as a 5-mm spherical region centered on the individual’s activation peak coordinate in a robust finger-tapping task (analyzed with FMRI Expert Analysis Tool). <u>Calculating each placement configuration took about 2 minutes, 3844 configurations together took about 128 hours.</u>
4. We found the individual’s optimal position *s*\* and orientation parameter *φ*\* as those with maximum average E-field strength in the ROI.
5. We calculated the individualized targeting effect map and optimal placement parameter for 8 participants (5 males, 3 females, mean age 26.4±2.1 years), and then summarized the normalized individualized targeting effect map into a group-level average targeting effect map and a standard deviation map. The group-level appropriate placement was the position and orientation that had a relatively large average targeting effect and relatively small standard deviation across participants.

**Figure 4.**
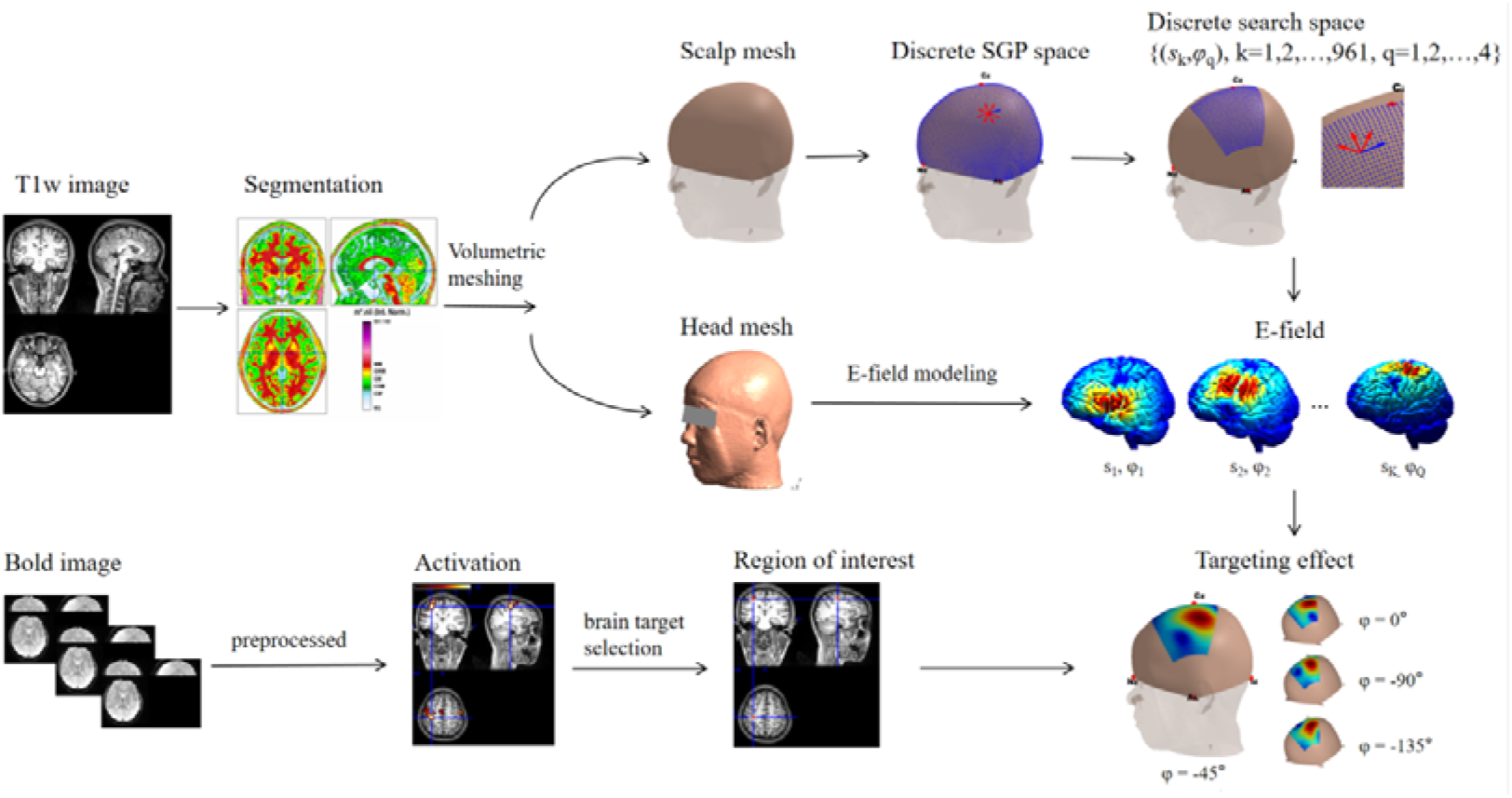
The SGP-based individualized optimization protocol

## 3. Results

### 3.1 Manual measurement experiment: evaluating speed, precision and reliability

**Table 1.** The speed, precision and reliability of SGP manual measurement.

| Speed | Mean | SD | Maximum | Minimum |
| --- | --- | --- | --- | --- |
| Measurement of the Cz point and the reference curve (step 1, 2) | (minute) |  |  |  |
| Technician 1 | 3.22 | 0.98 | 4.99 | 1.67 |
| Technician 2 | 3.64 | 0.42 | 4.70 | 3.24 |
| Technician 3 | 3.38 | 0.75 | 4.88 | 2.24 |
| Measurement of scalp position s (step 3) | (minute) |  |  |  |
| Technician 1 | 1.33 | 0.47 | 3.30 | 0.68 |
| Technician 2 | 0.91 | 0.24 | 1.62 | 0.51 |
| Technician 3 | 0.86 | 0.20 | 1.61 | 0.54 |
| Measurement of orientation $\phi$ with protractor (step 4) | (minute) | | | |
| Technician 1 | 0.59 | 0.22 | 1.13 | 0.37 |
| Technician 2 | 0.75 | 0.16 | 0.91 | 0.53 |
| Technician 3 | 0.57 | 0.23 | 0.80 | 0.36 |
| Precision | Mean | SD | Maximum | Minimum |
| Measurement of scalp position s | (mm) |  |  |  |
| Technician 1 | 3.74 | 2.15 | 10.94 | 0.95 |
| Technician 2 | 3.96 | 2.56 | 10.07 | 0.67 |
| Technician 3 | 4.60 | 2.74 | 11.57 | 0.65 |
| Measurement of 45° with protractor | (degree) |  |  |  |
| Technician 1 | 3.51 | 1.70 | 6.72 | 0.98 |
| Technician 2 | 6.38 | 1.76 | 9.18 | 4.25 |
| Technician 3 | 2.96 | 1.66 | 5.22 | 1.39 |
| Measurement of 45° with visual estimation | (degree) |  |  |  |
| Technician 1 | 4.50 | 2.02 | 8.06 | 1.93 |
| Technician 2 | 9.38 | 3.11 | 13.97 | 6.86 |
| Technician 3 | 7.65 | 3.54 | 13.28 | 3.30 |
| Measurement of other orientations with protractor | (degree) |  |  |  |
| Technician 1 | 4.36 | 3.10 | 6.31 | 0.78 |
| Technician 2 | 6.74 | 2.03 | 8.74 | 4.69 |
| Technician 3 | 2.91 | 1.46 | 4.30 | 1.38 |
| Reliability | Mean | SD |  |  |
| Intra-technician reliability | (mm) |  |  |  |
| SGP | 3.53 | 1.86 |  |  |
| Beam F3 (Trap et al.,2020) | 9.30 | 6.20 |  |  |
| 5.5 cm (Trap et al.,2020) | 11.70 | 6.80 |  |  |
| Inter-technician reliability | (mm) |  |  |  |
| SGP | 4.72 | 2.20 |  |  |
| Beam F3 (Trap et al.,2020) | 9.50 | 6.10 |  |  |
| 5.5 cm (Trap et al.,2020) | 12.20 | 6.60 |  |  |

Summing together the mean measurement time for each step in the placement process, the whole placement process (Step1,2 + Step3 + Step4) takes about 5.08 minutes in total. The longest is about 7.78 minutes and the shortest is only 3.12 minutes. The overall average accuracy of all technicians measured at all locations (9 locations) was 4.1±2.51mm. Among them, 73% of measurements’ errors were less than 5mm. The overall angle accuracy of all technicians at all angles (0°,45°,90°,135°) with protractor was 4.24±2.24°. We compared the measurement by protractor to visual estimation at 45° and found that errors from protractor were significantly smaller than errors from estimation (mean = 4.09° versus 6.51°, t = 2.9668, df = 46, *p*<0.01), but using a protractor required more time. For both intra-technician and inter-technician reliability, the SGP-based protocol (intra-technician: 3.53±1.86mm; inter-technician: 4.72±2.2mm) performed better than the BeamF3 method (intra-technician: 9.3±6.2mm; inter-technician: 9.5±6.1mm) and 5.5cm method (intra-technician: 11.7±6.8 mm; inter-technician: 12.2±6.6 mm), as reported in Trapp et al., 2020.

### 3.2 Demonstration experiment for individualized and group-based optimization

TMS electrical effects in the motor cortex were modeled for 961 positions and 4 orientations. As shown in Fig. 5A, both different positions and orientations will lead to different electrical effects at the ROI. The personalized optimal positions were (0.52,0.36), (0.54,0.34), (0.56,0.22), (0.52,0.36), (0.46,0.46), (0.52,0.38), (0.52,0.36) and (0.46,0.36) for the seven participants. These positions were distributed around the 10-20 landmark C3 (0.50,0.35). The individually optimal orientations included all searched orientations, 0°, -45°, -90° and -135°, showing inter-individual variation in orientation. The group-based appropriate parameter was (0.52,0.36, -90°), with a maximum average targeting effect *μ* = 0.83 and a relatively small standard deviation *σ* = 0.14.

**Figure 5.**
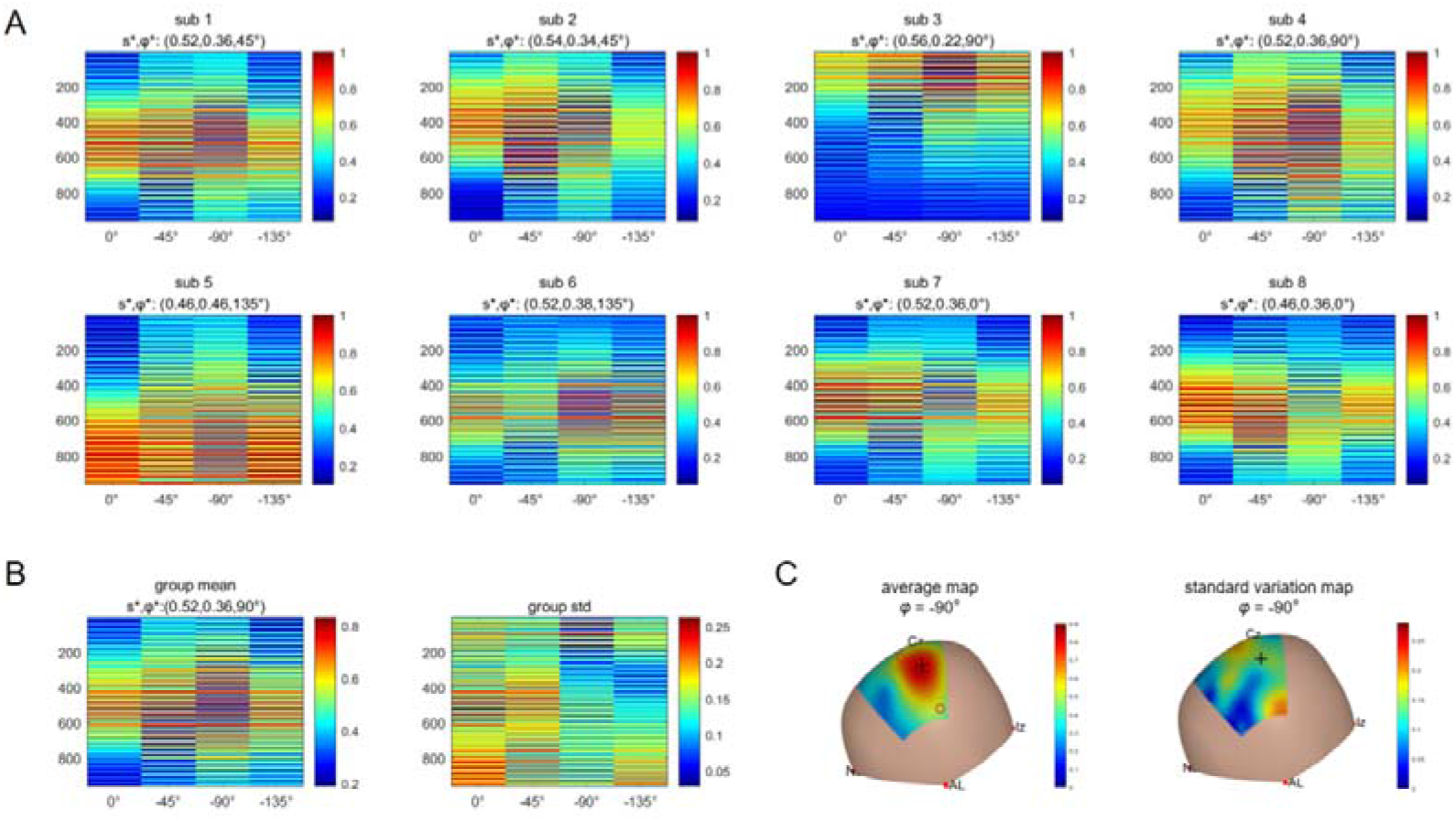
A) Individualized targeting effects for 961 positions (y-axis) and 4 orientations (x-axis). Optimal position and orientation (*s*\*, *φ*\*) that maximizes individualized targeting effect is shown in the subtitle. B) Group-based targeting effect map, standard deviation map and the group-appropriate parameters. C) Scalp visualization of group-based targeting effect map and the standard deviation map at group-appropriate orientation (*φ* = -90°). Individualized optimal positions that maximize individual targeting effects at *φ* = -90° are marked with red circles; the group-based appropriate position is marked with a black cross.

## 4. Discussion

In this article, we proposed a scalp-geometry based parameter-space to describe TMS coil placement and showed how to apply SGP in different clinical settings for coil placement optimization and implementation.

### 4.1 Properties of scalp geometry-based parameter

SGP describes the position and orientation of TMS coils based on the geometric properties of the head. For the coil position *s*, the geometry information to be measured includes the position of five reference points (NZ, IZ, CZ, AL, AR) and the arc length of four geodesics (NZ-*s*’, NZ-IZ, AL-*s* and AL-AR). For the orientation of the coil at point *s*, the geometric information to be measured includes the intersection line of the tangent plane at point *s* and the plane of the active arc, and the rotation angle of the line around point *s*.

This scalp geometry-based definition leads to several basic properties. First, a set of parameters in SGP space has a one-to-one correspondence to a conventional TMS coil placement. Given a conventional coil placement on the scalp, each coil center location corresponds to a unique position parameter *s* defined by two arc length proportions (P_NZ_,P_AL_) and each coil handle direction corresponds to a unique rotation *φ* from the zero vector of *s* in the tangent plane. Conversely, for any given set of SGP parameters (*s*, *φ*), there is a unique corresponding coil placement: placing the coil center at s and turning the handle *φ* degrees from the zero vector in the tangent plane. This one-to-one correspondence allows SGP-based methods to accurately navigate and record any possible scalp position and orientation.

Second, the parameter space has inter-individual consistency. That is, if we place coils on different individuals’ scalps using the same SGP parameter, these placements are consistent. That is a natural property shared by both the 10-20 system and the SGP. They use reference points to define subsequent curves and measure proportional positions on the curves. Reference points are scalp landmarks consistent between different people and the proportional measurement take head size variations into account. Therefore, the proportional position *s*’ on the reference curve has inter-individual consistency, and the active curve defined by AL, AR and *s*’ also has consistency. The direction of the intersection between the active curve plane and the tangent plane of *s* is consistent, thus the coil handle direction a certain angle from this intersection direction also has inter-individual consistency. These properties allow the SGP to be used as a standard space to compare and synthesize results from different individuals or different studies, to obtain more generalizable conclusions. As shown in Fig. 5, preliminary results show that the differences in targeting effect distributions among different individuals are not very large (std<0.27 normalized effect) so there are similarities that support synthesis.

Third, the SGP is quantitatively measurable both *in silico* and *in vivo*. All the information needed to define SGP are measurable geometric parameters on the head, which provides the theoretical basis for measurability. *In silico*, we can reconstruct the scalp mesh from MRI images, calculate arbitrary SGP-based positions and orientations and use computational modeling such as electric field simulation to traverse over the targeting effects of potential placements (as shown in the demonstration experiment). *In vivo*, the scalp mesh can be reconstructed from a dense sampling of a participant’s scalp with a 3D digitizer. Any error between the current placement’s SGP position and the target one can be computed in real-time to help achieve accurate coil placement. Meanwhile, positions in SGP can be measured manually in clinical practice, in a fast and simple way that is free of additional equipment. As shown in the methods, the measurement only involves measuring four geodesic distances: NZ-*s*’, NZ-IZ, AL-*s* and AL-AR and one angle from the zero direction. In our manual measurement experiment, we recorded the speed, precision and reliability of three technicians while they performed these steps. The speed of placement (about 5 minutes) was quick compared to the reported time required for the 10-20 method to only measure the position (16 minutes, Xiao 2016). Our error was <5mm in 73% of measurements, comparable to the reported discrepancies of the BeamF3 (<9.9 mm in 75% of participants). Our reliability (intra-technician: 3.53±1.86mm; inter-technician: 4.72±2.20mm) is better than the BeamF3 method (intra-technician: 9.3±.6.2mm; inter-technician: 9.5±6.1mm) or 5.5cm method (intra-technician:11.7±6.8 mm, inter-technician:12.2±6.6 mm) for both intra-technician and inter-technician reliability (Trapp et al., 2020). These experimental results show that, in the absence of navigational equipment and complex computing capacity, coil parameters can be manually measured on the scalp with good accuracy and speed.

### 4.2 Comparison of SGP with 10-20 and CDP methods

How to describe the position and orientation of coil placement is the common basis of optimization and actual placement. Different description methods for coil placement may have different effects on optimization and actual placement. Below, we compare our description method with two existing description methods from three perspectives: individualized optimization, group synthesis and real-world implementation of the coil placement protocol.

#### Performance of different methods in individualized optimization

The goal of optimization is to go through all possible coil placements to find the best placement. Therefore, it is important for a description method to be able to describe all potential coil placements. Coil placements can be divided into physically unachievable (part of the coil intersects with the head) and physically achievable placements. Physically achievable placements also include large areas far away from the scalp that are not able to generate efficient stimulation due to electric field attenuation. Therefore, the physically realizable space with practical significance is mainly composed of conventional placements (coil placed in a plane tangent to the scalp with coil center contacting the scalp). Conventional placement space contains the commonly used placements in clinical practice and is adopted in most optimization studies as the search space.

The 10-20 method and its extensions (like 10-10 an 10-5) are essentially a set of finite points rather than a continuous parameter space. It can only describe a limited number of protocols in conventional placement space, which is a significant problem for optimization. On the other hand, CDP is a continuous space that can quantitatively describe position and posture of any coil placement, which is conductive to global optimization. However, taking the space occupied by the person’s head into account, many parameters in CDP are physically impossible or far away from the head. Therefore, CDP-based optimization usually requires parameters to be constrained to the conventional placement space, which leads to a complex constraint optimization problem. The proposed SGP space is based on head geometry. All SGP parameters are constructed according to head geometry, therefore it only includes conventional placement with coil center contacting the scalp tangentially, removing any intersecting or far away placements, which reduce the size of the search space and improve the efficiency of optimization. Moreover, optimization based on SGP is an unconstrained optimization problem, which is usually much simpler than constrained optimization problems in a mathematical sense. Meanwhile, SGP is also a continuous space, so it will not leave out any possible conventional placement, as compared to 10-20 methods and their extensions.

#### Performance of different methods in group synthesis

As mentioned in the Introduction, in some clinical practice individualized optimization is impossible due to the absence of MRI scans or lack of time and computing power. Synthesis of individualized results from previously recorded databases into a group atlas can be an important alternative to guide placement in such cases. Inter-individual consistency of coil placement is a requisite for synthesis. Although the descriptive capacity of the 10-20 system and its extensions are insufficient, 10-20 system points have natural correspondence between individuals, which enables the optimization results based on 10-20 system to be summarized directly. For example, Gomez-Tames (2019) synthesize individual deep TMS dosage in targeted deep brain regions at different 10-20 positions into a group-level dosage to overcome the limitations of using individualized head models to characterize coil performance in a population, providing a group-level dosage atlas to guide coil placement for each deep region. Like the 10-20 system, the definition of SGP is also determined based on the geometric parameters of the head, which are consistent between individuals. Therefore, the optimization results from different individuals can be summarized directly, as shown in our demonstration experiment.

CDP parameters, on the other hand, are based on individual-specific three-dimensional coordinate systems: the space where individualized electric field modeling is performed, i.e., the native T1 image space. So, in order to synthesize the result of different individuals, their CDP parameters in native space have to be registered to a standard CDP space based on MNI space. That means an additional registration process is needed. Although the registration between spaces (via images) based on brain intensity values or brain surface geometries is commonly done in many software, there is much less work on how to perform a registration between scalps, where the coil position is located. Moreover, this process not only involves the registration of coil position points, but also involves the registration of a directional vector, which is even more complicated. Little effort has been done in this area. Another feasible way is to convert the optimization result from individual CDP space into SGP space (each three-dimensional coordinate can be converted to a proportional coordinate *s* and each set of three three-dimensional vectors can be convert to a rotation angle *φ* in the tangent plane of *s*), then optimized results can be summarized directly.

#### Performance of different methods in real-world implementation

The actual stimulation effect of TMS depends not only on the computational optimization of coil placement parameters, but also on whether the coil can be placed accurately according to the optimal parameters on a person’s head *in vivo*. Ideally, precise placement can be achieved with a neuronavigation system. However, expensive neuronavigation systems are not always available. If manual measurement can be done quickly and simply while sacrificing little accuracy, it can be practical for clinical settings. According to the definition, each 10-20 point can be measured manually by a sequence of steps, with later measurements dependent on prior measurements. However, multiple measurements and calculations can be excessively time-consuming (e.g., measuring the P4 point can take about 16 minutes as reported in Xiao et al., 2016). In addition, with more measurements comes more opportunity for human error. BeamF3 is a simpler and faster method that was proposed to improve the speed of finding the F3 position for prefrontal coil placement. It simplified the F3 position measurement steps to only three skull measurements. However, its transformation formula only works for F3 and cannot be generalized directly to other locations. Mir-Moghtadaei has published a series of articles (Mir-Moghtadaei et al., 2015, 2016, 2017) to establish a scalp heuristic that can transform CDP positions to skull measurement (e.g., 25% Nasion-Inion for locating the dorsomedial prefrontal cortex). It is more precise than the BeamF3 method, as it is based on individuals’ MRI scans. However, in cases where MRI scans are not available, the scalp-based heuristic needs an anatomical MRI database and cannot be generalized to other locations.

The SGP has obvious advantages in coil placement implementation. In cases where an SGP-based navigation system (navigate to *s* and *φ* directly on the scalp, in development) is available, both the individual and group optimal placement can be implemented accurately. For those who do not have an SGP navigation system but have traditional CDP-based neuronavigation systems like BrainSight, we provide an alternative. That is, if the individual’s T1 image is available, the SGP individual parameters can be transformed into CDP parameters and used by BrainSight (using the T1 image to sample the scalp mesh and calculate the CDP position and direction vectors corresponding to (*s*, *φ*) on the scalp mesh, then using an NDI optical camera to register the mesh in computational space and the person’s scalp in the real world, then navigating to the CDP position and direction vectors). In the absence of navigation systems, SGP parameters can also be implemented by manual measurement of *s* and *φ*. Each scalp placement can be measured independently, therefore providing a very fast (< 5 minute), accurate (73%<5 mm error) and low-cost protocol to realize arbitrary coil placement, which can be especially useful in large-throughput clinical practice.

Taken together, the SGP method combines the advantages of the 10-20 method and the CDP method in individual optimization, group synthesis and real-world implementation. When considering these three aspects simultaneously, SGP provides an improved coil placement protocol that can be flexibly applied to different situations.

### 4.3 Limitation and further directions

#### Extension of scalp geometry-based parameters

The limitations of the SGP are related to its advantages, that is, the SGP is constrained to conventional placements on the scalp. Although conventional placement represents the main coil placement protocol in clinical practice and most optimization studies use conventional placement space as the search space, conventional placement does not include all physically achievable coil placements. For example, the SGP-based parameters do not include a tilt angle from the tangent plane. The hand-held coil placement cannot always be strictly located on the tangent plane and will thus introduce tilt error. Some studies have pointed out that the tilt angle will affect the electric field distribution. So, it may be useful to record the tilt angle for hand-held placement for more accurate modeling and understanding of TMS effects. However, it is difficult to record such an angle without the help of a navigation system, so this remains a practical problem.

Another problem is that the current discrete SGP is established by uniform interval sampling of the *s* and *φ* parameters. The uniform sampling of *s* (continuous proportional coordinate) can result in nonuniform Euclidean distances between unit-value coordinates. As shown in Fig. 1C, sampling points are sparse around the vertex (inter-point Euclidean distance of about 3.7mm), while much denser around the ears (inter-point Euclidean distance can reach less than 1 mm). If the search space is not near the ears and is in a small area, discrete SGP can be approximately uniform in Euclidean distance; but if the search space is the full head, the equal interval SGP parameter sampling may not be appropriate as it will generate too many points around the ear. An improved discrete sampling based on the Fibonacci lattice method (González, 2010) can help to make the resulting Euclidean distances uniform.

#### Extension of SGP-based coil placement optimization

Our preliminary results showed inter-individual differences in optimal placement parameters. This may be a true individual difference: like Balslev (D et al., 2007) reported a range of 63° in optimal orientations across different participants. However, it may depend on the selection of ROI: even in the motor area, the most common optimal orientation varies when targeting different precise locations (Gomez-Tames et al., 2018b). It may also depend on the selection of E-field variables: we used the E-field magnitude (which leads to the same result for *φ* and 180°-*φ*); using the perpendicular component and tangential component may lead to significantly different optimal coil placements (Gomez et al., 2021). Our demonstration mainly follows the protocol proposed by Balderston (2020), that chose to maximize the E-field magnitude in the 5-mm ROI centered at the functional activation peak. However, the model for calculating simulated effects needs further refinement, and further studies can explore other optimization strategies to provide a more comprehensive understanding.

In addition to optimizing the primary TMS effect in the targeted ROI via electric modeling, we can also optimize based on secondary network effects propagated to distal regions via functional connectivity. For example, we can model anti-depressant effects by calculating the resting-state connectivity between DLPFC and subgACC (Fox et al., 2012; Siddiqi et al., 2020), where more anti-correlated connectivity leads to better efficacy. The electric modeling and functional connectivity modeling can also be combined into one optimization protocol (Balderston et al., 2020).

Besides using computational modeling effects to optimize placement, we can also use TMS clinical effects for optimization by continuously recording TMS coil placement and treatment effects in clinical studies and performing a meta-analysis to find placements with better effects. One problem that hinders the synthesis is the inconsistency of recorded descriptions. As reviewed in Lefaucheur et al., 2020, some records use 10-20 descriptions like F3 to report the original scalp location, while others report MNI coordinates, which are line projection results from the original scalp position in CDP space. The SGP is a unified system compatible with different descriptions. It can summarize existing conventional coil placements recorded in various manners by converting them into SGP parameters, regardless of whether the coil positions were described using 10-20 method, 5-cm method or the targeted brain locations obtained from a line projection. Therefore, SGP can provide a standard framework to facilitate synthesis of TMS effects at different positions and orientations from different studies. SGP can be used to map efficacies of different kinds of TMS treatments, like anti-depressant effects, motor evoked potential effects and so on, resulting in disorder-specific coil placement atlases to guide clinical practice.

Finally, in addition to describing and optimizing TMS coil placement, SGP is also suitable for other transcranial techniques that place optodes or electrodes on the scalp. For example, SGP can be used to describe and optimize the electrode positions for transcranial direct current stimulation (tDCS).

## Acknowledgements

This work was supported by the National Natural Science Foundation of China (grant no. 82071999 and 61431002). The authors declare no competing financial interests. We thank Ye Xin for the assistance in conducting the experiments.

## Notes

### Competing Interest Statement

The authors have declared no competing interest.

